# Cross-taxon Coarse-Grained IDP Simulations Enable Architecture-Independent Generalization

**DOI:** 10.64898/2026.08.18.745604

**Authors:** Juan Velasquez, Taseef Rahman

## Abstract

Intrinsically disordered proteins and regions are found across all kingdoms of life, yet the computational characterisation of their conformational ensembles has remained almost entirely confined to the human proteome. Whether the force fields used to generate them remain accurate for taxonomically distant organisms, and whether the sequence-ensemble relationships they reveal transfer across taxa well enough to improve prediction on phylogenetically held-out organisms, are open questions. Here we introduce BENDER, a dataset of 11,533 IDP sequences spanning 13 taxonomic groups, each simulated under CALVADOS-2 molecular dynamics and annotated with ensemble-level geometric and novel contact-network properties, together with per-sequence pi–pi and cation–pi contact frequencies linked to phase-separation propensity. We show that CALVADOS-2 ensembles agree strongly with an orthogonal structural reference across the full dataset, with both held-out taxa performing above the dataset median, and that direct comparison against a second independently parameterized force field reveals no systematic scaling-exponent bias. We find that cross-taxon training data improves out-of-distribution ensemble prediction in two independent architectures despite training on one-third the data. Positive degree assortativity is conserved across all taxonomic groups, suggesting that hub topology in disordered protein contact networks is a conserved physical feature of sequence-encoded disorder rather than an evolutionary contingency.

## 1 Introduction

Intrinsically disordered proteins and regions (IDPs and IDRs) constitute a substantial fraction of proteomes across all domains of life, from the signalling and regulatory networks of mammals to the stress-response systems of bacteria and the replication machinery of RNA viruses [1, 2, 3]. Unlike folded proteins, IDPs do not adopt a stable three-dimensional conformation but instead exist as heterogeneous ensembles of rapidly interconverting structures whose conformational properties underlie their functional versatility. This structural plasticity enables IDPs to perform central roles in signalling, transcriptional regulation, biomolecular condensate formation, and viral replication [4, 5], and their dysregulation is implicated in neurodegeneration, cancer, and infectious disease [6, 7]. Despite this broad taxonomic reach, the computational characterisation of IDP conformational ensembles has remained almost entirely confined to the human proteome [8, 9, 10, 11, 12].

The dominant resource for IDP ensemble prediction is Human-IDRome [9], a dataset of 28,058 human intrinsically disordered regions with conformational ensembles generated by the CALVADOS-2 coarse-grained force field [13]. Building on this foundation, GeoGraph [8], along with ALBATROSS [10] and STARLING [11] (using different datasets) have demonstrated that ensemble-averaged properties such as the radius of gyration and polymer scaling exponent can be predicted from sequence alone at proteome scale. Yet this progress rests on a narrow evolutionary foundation, as Bacteria, Archaea, Fungi, Viruses, Protists, and Plants all harbour IDPs with distinct compositional patterns, evolutionary pressures, and functional roles absent from human-centric datasets [2, 4, 5], and evaluation has been performed exclusively on the same human-centric distribution used for training, meaning that failure on non-human sequences would have gone undetected. Indeed, a recent study on generative IDR design independently identified the absence of large, systematically annotated cross-taxon datasets as the primary bottleneck for the next generation of data-driven methods [14].

This taxonomic confinement leaves two questions open that are fundamental to the simulation community. The first is whether CALVADOS-2, parameterized against SAXS and NMR data from eukaryotic IDPs [13], produces reliable ensembles for taxonomically distant organisms such as bacteria, archaea, and viruses, groups for which no cross-force-field validation at proteome scale has been conducted. The second is whether the sequence–ensemble relationships discovered in human IDPs, including polymer scaling behavior, chain compaction patterns, and contact-network topology, represent conserved physical laws that apply wherever disordered proteins are found, or artifacts of the evolutionary pressures that shaped the human proteome. Whether these relationships are universal determines both whether current predictors can be trusted on non-human sequences and whether the physical principles they encode carry genuine evolutionary generality.

Here we introduce BENDER (**B**iological **E**nsembles of **N**atively **D**isord**E**red p**R**oteome), the first cross-taxon IDP ensemble dataset, comprising 11,533 sequences spanning 13 taxonomic groups simulated under CALVADOS-2 molecular dynamics. Beyond the geometric properties characterized in previous datasets, BENDER introduces novel contact-network descriptors—specifically, global efficiency, fragmentation index, average clustering coefficient, transitivity, and degree assortativity—along with per-sequence pi–pi and cation–pi contact frequencies linked to phase-separation propensity [15]. To quantify the contribution of training distribution to out-of-distribution performance, we additionally introduce KESTREL, a lightweight transformer trained on BENDER sequences, alongside KESTREL-Human, an architecture-matched variant trained on Human-IDRome for a controlled comparison.

These experiments address both questions raised above. The reliability of CALVADOS-2 ensembles on held-out Viruses confirmed against an orthogonal structural reference and a second independently parameterized force field indicates that the core physics governing disordered chain compaction is not an artifact of the eukaryotic sequences used in force-field parameterization, opening the way for routine cross-taxon IDP simulation without separate per-group validation. The consistent improvement from cross-taxon training data, observed across two architecturally distinct models, reveals that sequence-ensemble relationships appear substantially conserved across deep evolutionary time, and that the human proteome represents a narrower window of accessible sequence space than its dominant position in the IDP training literature implies. The conservation of positive degree assortativity across all 13 taxonomic groups, combined with the high predictability of contact-network global efficiency from sequence alone on both held-out taxa, suggests that distributed hub topology in disordered protein contact networks is a conserved physical feature rather than an evolutionary contingency of any particular lineage.

We make the BENDER dataset available at https://huggingface.co/datasets/taseef/BENDER, and the KESTREL model code, weights, and all analysis scripts along with per-sequence target values are publicly released at https://anonymous.4open.science/r/IDP-Project-17EA.

## 2 Related Work

Coarse-grained molecular dynamics has made proteome-scale IDP ensemble generation feasible. CALVADOS-2 [13] is a one-bead-per-residue model parameterised against experimental SAXS and NMR measurements of eukaryotic IDPs, and forms the basis of Human-IDRome [9], the largest existing IDP ensemble dataset comprising 28,058 human disordered regions. MPIPI-GG [16] uses independently derived interaction potentials with no shared parameterisation data, as do other coarse-grained frameworks including HPS [17] and ABSINTH [18]. Despite the availability of multiple force fields, no systematic cross-force-field validation of IDP ensemble properties has been conducted at proteome scale, and the reliability of any coarse-grained force field on non-eukaryotic sequences has not been tested.

Human-IDRome has catalysed a series of supervised predictors, among which ALBATROSS [10] predicts geometric ensemble properties using a BiLSTM trained on MPIPI-GG trajectories, and GeoGraph [8] combines ESM-2 embeddings with a graph neural network architecture informed by residue-level contact networks, trained on Human-IDRome with global efficiency as an auxiliary target. Several other approaches have applied protein language models, generative diffusion, and ensemble deep learning to IDP property prediction [11, 12, 19], though all existing datasets and benchmarks remain restricted to human or eukaryote-centric sequences, and all existing simulation datasets model IDR fragments extracted from multidomain proteins, introducing domain context effects that systematically alter conformational behaviour [20].

Single-chain IDP ensemble properties are experimentally established as connected to multi-chain phase behaviour. Dignon et al. [17] demonstrated a strong correlation between single-chain collapse temperature and phase-separation critical temperature. Martin et al. [21] showed that aromatic residue valence determines both single-chain compaction and the full phase diagram for prion-like domains, and Bremer et al. [22] confirmed this coupling for naturally occurring sequences while identifying conditions under which it weakens. Vernon et al. [15] established pi–pi contact frequency as the primary sequence feature predicting phase-separation propensity. These connections motivate the inclusion of pi–pi and cation–pi contact frequencies in the BENDER dataset alongside conventional geometric targets.

BENDER addresses the cross-taxon gap directly, providing the first dataset of IDP ensembles spanning 13 taxonomic groups with force-field validation against both an orthogonal computational reference and a second independently parameterised force field.

## 3 The BENDER Dataset

### 3.1 Sequence Curation

Sequences were sourced from MobiDB[23] applying a consensus disorder filter requiring 100% disorder across the full sequence and a minimum length of 30 residues. Where the initial query for a taxonomic group was dominated by a single species, e.g. *Oryza sativa* in Plants and *Plasmodium falciparum* in Protists, additional sequences from taxonomically diverse species were incorporated to ensure representative taxon-level coverage. Taxonomic group representation reflects both the availability of fully disordered sequences in MobiDB[23] and the underlying biology: Bacteria and Plants contribute the largest numbers (2,850 and 2,480 respectively), consistent with the high abundance of standalone disordered proteins in bacterial stress response and plant regulatory networks [24, 25]; Mammals contribute fewer sequences (1,361) despite their well-characterized proteome, as human Disordered Regions frequently occur as short disordered motifs embedded within multi-domain proteins that fail the 100% disorder criterion applied here; and Viruses (1,025) and Protists (1,049) contribute comparable numbers, reflecting high per-proteome disorder content [2, 3] offset by smaller proteome sizes and sparser database coverage relative to cellular organisms.

To eliminate redundancy, global CD-HIT clustering at 40% sequence identity was applied across all taxonomic groups simultaneously, ensuring that cross-taxon universality claims cannot be attributed to shared ancestral sequences. Where a single species dominated the initial query for a taxonomic group, additional sequences from taxonomically diverse species were incorporated to ensure representative taxon-level coverage. The final dataset comprises 11,533 sequences drawn from 1,958 unique organisms spanning 13 taxonomic groups (Figure 1). A further species-level breakdown is provided in Supplementary Figure S1

**Figure 1:**
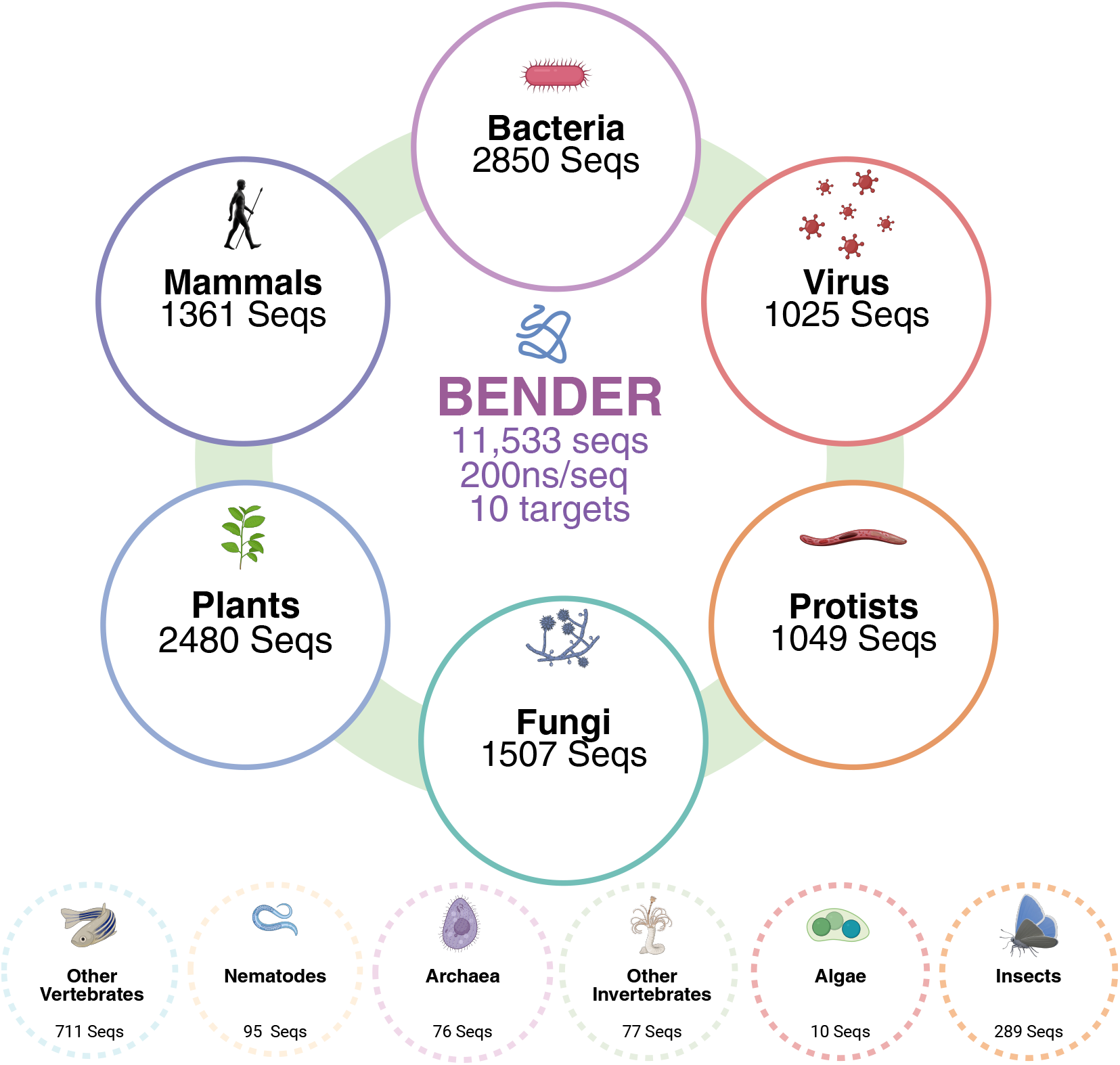
BENDER dataset showing 11,533 IDP sequences across 13 taxonomic groups with CALVADOS-2 conformational ensembles, 10 ensemble-level prediction targets, and per-sequence pi-pi and cation-pi contact frequencies.

### 3.2 Simulation Protocol

All simulations were performed using the CALVADOS-2 coarse-grained force field [13], which represents proteins at single-bead-per-residue resolution and has been extensively validated against SAXS, smFRET, and NMR experimental data for IDPs. Simulations were conducted in the NVT ensemble at 300 K with an ionic strength of 0.15 M NaCl. Every sequence was simulated for a minimum of 200 ns, with sequences exceeding 150 residues receiving extended simulation time scaled as *L*^1.5^ to account for longer reconfiguration timescales. The first 50% of each trajectory was discarded as equilibration, resulting in a minimum production phase of 100 ns per sequence. Simulations were initialised from extended-chain conformations, avoiding the initial-structure bias that affects AlphaFold-initialised IDR simulations [26, 27]. Generating the full dataset required approximately 750 GPU-hours of coarse-grained MD, with minimal overlap with Human-IDRome as only 39 human IDPs appear in both.

### 3.3 Prediction Targets

Ten ensemble-level prediction targets were computed for each sequence, spanning geometric and contact-network properties. Geometric targets include the radius of gyration (*R_g_*), end-to-end distance (*R_e_*), Flory scaling exponent (*ν*), asphericity (Δ), and Flory prefactor (*A*_0_), which are standard descriptors of IDP chain dimensions and shape.

Contact-network properties were computed from the ensemble-averaged intrachain contact graph (*C_α_*–*C_α_* cutoff 0.8 nm) and comprise global efficiency (*g*_eff_), fragmentation index, average clustering coefficient, transitivity, and degree assortativity. Global efficiency measures how effectively conformational signals propagate across the chain [28], while degree assortativity captures whether highly connected residues preferentially contact other highly connected residues, reflecting hub architecture implicated in condensate material properties [29, 30]. Per-taxon target distributions are further visualized in Supplementary Section I. These constitute a qualitatively new class of IDP descriptors unavailable in any existing dataset, and no prior model predicts them from sequence.

Per-sequence pi–pi and cation–pi contact frequencies are computed from the simulation trajectories and included as targets linked to phase-separation propensity [15], extending condensate-relevant sequence features to all 13 taxonomic groups.

### 3.4 Force-Field Validation

The central methodological question for any simulation-derived dataset is whether the labels reflect physical behaviour or force-field-specific artefacts. This question is sharpened for BENDER because CALVADOS-2 was parameterised against eukaryotic experimental data, yet the dataset includes sequences from all three domains of life. We provide three independent lines of validation.

CALVADOS-2 is parameterised and validated directly against experimental SAXS and NMR [13], and Human-IDRome confirmed this accuracy against experimental SAXS at dataset scale [9], establishing an experimental foundation for the force field’s labels. Our first validation test examines whether this accuracy holds on the taxonomically distant sequences that constitute the majority of BENDER, using AlphaFold2 PAE as an orthogonal computational reference whose uncertainty estimates derive from evolutionary co-variation rather than any physical force field. Following Tesei et al. [9], who applied this to 4 proteins, we compute per-sequence Pearson *r* between inter-residue distance standard deviations from each CALVADOS-2 trajectory and the corresponding AF2 PAE matrix, a roughly 3,000-fold increase in coverage. Across the full dataset (*n* = 11,533), the median correlation is *r* = 0.804, with OOD taxon Viruses at 0.819 (Figure 2a).

**Figure 2:**
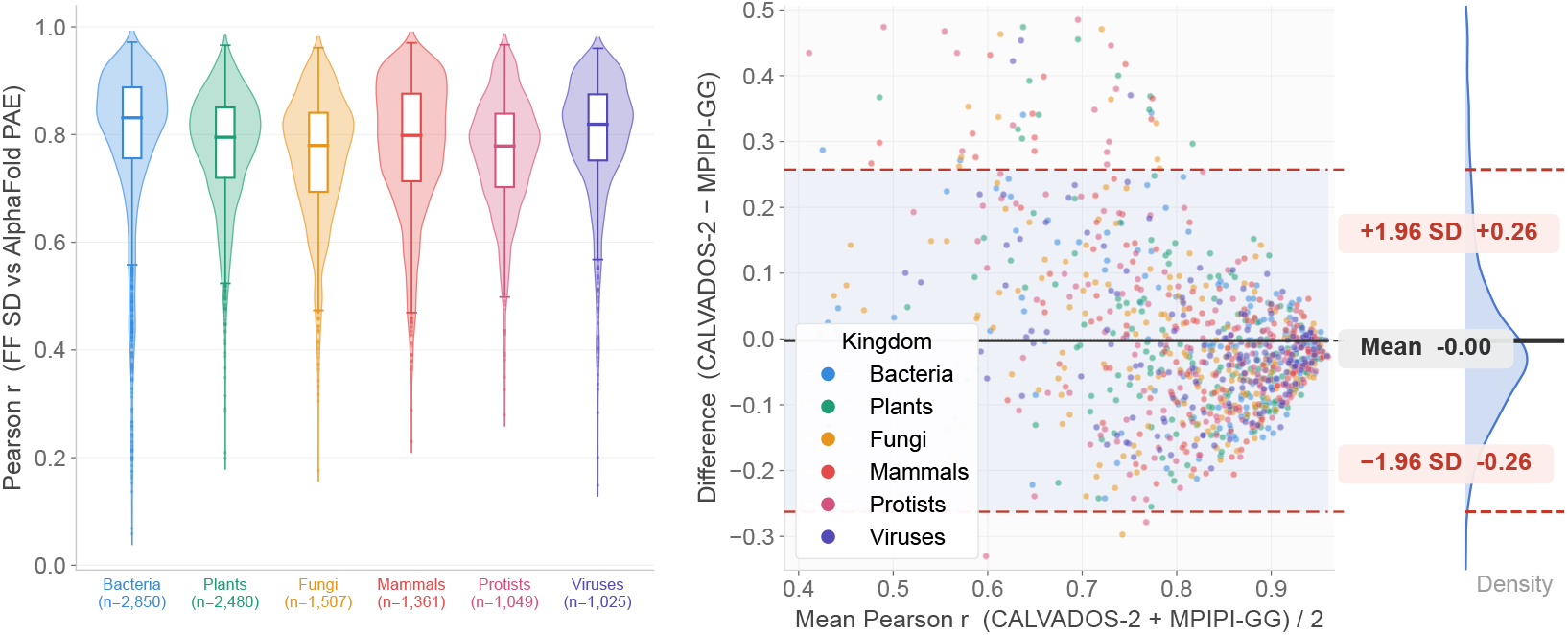
**(a)** Per-sequence correlation between force-field trajectory SD and AlphaFold2 PAE, shown per taxon for the 992 paired sequences. Both force fields achieve median *r >* 0.8 across all six kingdoms. **(b)** Bland-Altman plot comparing the two force fields’ agreement with AF2 PAE on a per-sequence basis. Mean difference −0.003 (solid), 95% limits of agreement [−0.263, +0.257] (dashed). The near-zero mean confirms that neither force field systematically agrees better with AF2 PAE than the other, and the spread reflects per-sequence variability rather than systematic bias.

**Figure 3:**
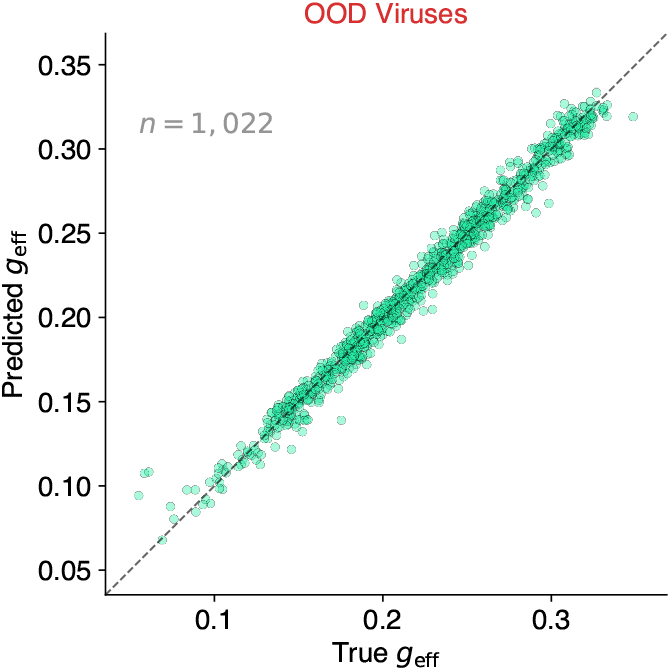
Predicted vs true *g*_eff_ for 1,022 OOD viral sequences (*R*^2^ = 0.981). No prior model predicts this target.

For the cross-force-field comparison, we simulated 992 of the BENDER sequences under MPIPI-GG [16], a force field with different interaction potentials sharing no parameterisation data with CALVADOS-2. Both were validated against AF2 PAE on the same subset. CALVADOS-2 achieves median *r* = 0.830 and MPIPI-GG 0.868, with both rising on the 163 viral sequences to 0.850 and 0.880. To our knowledge, this is the first validation of MPIPI-GG against an orthogonal reference. Direct *ν* comparison shows no systematic bias, with a pooled *r* = 0.698, mean Δ*ν* = −0.0004 (SD 0.045, 95% LoA [−0.089, +0.088]), and OOD Viruses showing the highest per-kingdom agreement (*r* = 0.777, Figure 2b). A variance decomposition attributes approximately 20% of the disagreement to pipeline noise and approximately 80% to genuine force-field differences. We do not explain this away as noise, as the force fields differ physically on *ν*, and we state this as a limitation.

On 32 sequences shared with Human-IDRome, *ν* values agree at bias +0.00004 (95% CI [−0.006, +0.006], MAE 0.015). Flory fits are well determined, with a median fit *R*^2^ = 0.998 (95.2% above 0.95) and median *ν* uncertainty 0.0025 (3.9% of population SD). Convergence quality is consistent across taxa, with all groups maintaining median fit *R*^2^ above 0.99 (Figure 4) and the small proportion of sequences below the threshold distributed across groups without systematic bias.

**Figure 4:**
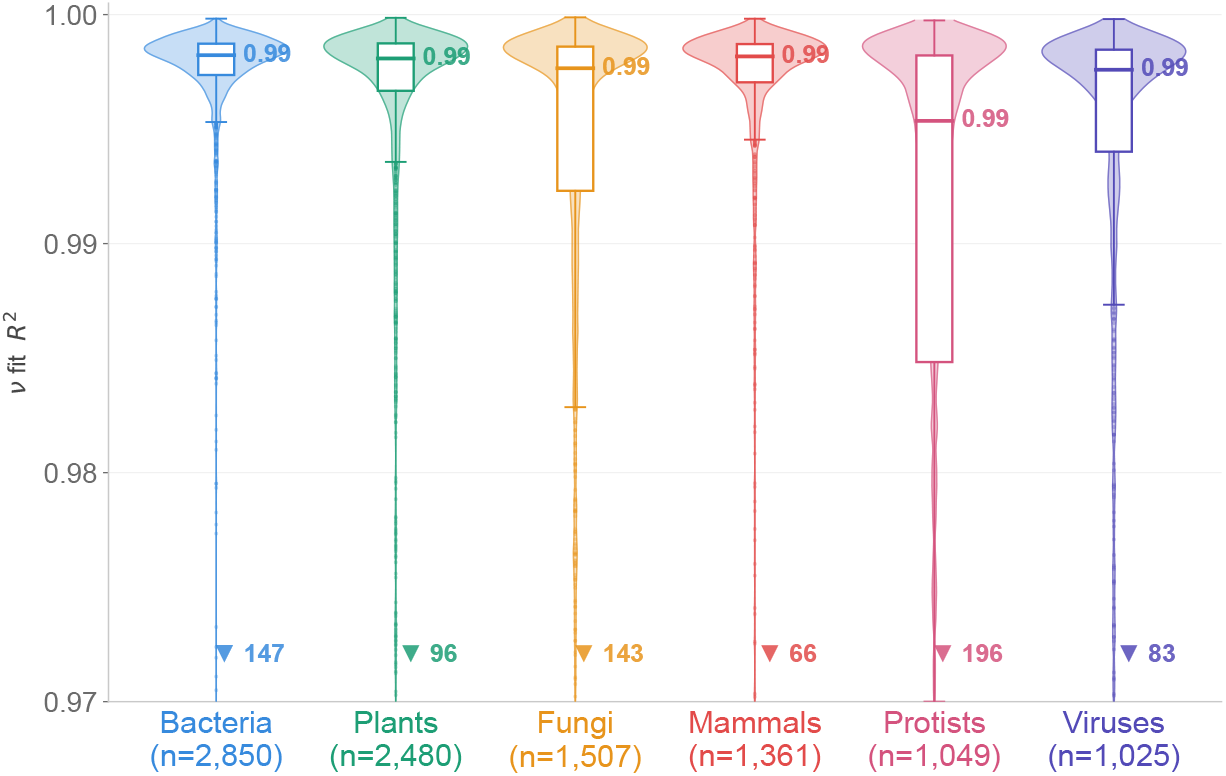
Per-taxon Flory scaling fit *R*^2^. All taxa maintain median above 0.99. Numbers below each violin indicate sequences with fit *R*^2^ < 0.95. Axis truncated at 0.95 for visibility.

## 4 Results

To determine how training distribution shapes out-of-distribution generalisation in IDP ensemble prediction, we designed a controlled comparison in which architecture and training data are varied independently. KESTREL is a 2.2M-parameter transformer trained from scratch on one-hot encoded BENDER sequences, with KESTREL-Human as an architecture-matched counterpart trained on Human-IDRome [9], isolating the effect of training distribution within a single model family. GeoGraph [8], a graph neural network trained on Human-IDRome, is paired with GeoGraph-BENDER, the same architecture retrained on BENDER, isolating the same question in a second, fully distinct architecture. PhyschemMLP, a 20k-parameter compositional baseline with no positional information trained on both datasets, serves as a lower bound on sequence-based prediction. Full architectural and training details are in the supplement, and all results are mean ± SD across 3 seeds.

We exclude ALBATROSS [10] from the benchmark table because three documented confounds make its *ν* and *A*_0_ entries non-comparable with those of the other models. These include a Flory scaling convention difference, a difference in *R*^2^ definition between squared Pearson correlation and coefficient of determination (discussed further in Quinn et al. [8]), and a potential force-field mismatch arising with validating ALBATROSS (which is trained on MPIPI-GG trajectories) against CALVADOS-2 labels. Subsequently, we find that the absolute *ν* error of MPIPI-GG against CALVADOS-2 is flat across all taxa (MAE 0.022–0.034), confirming the confound is systematic rather than taxon-dependent.

**Table 1:**
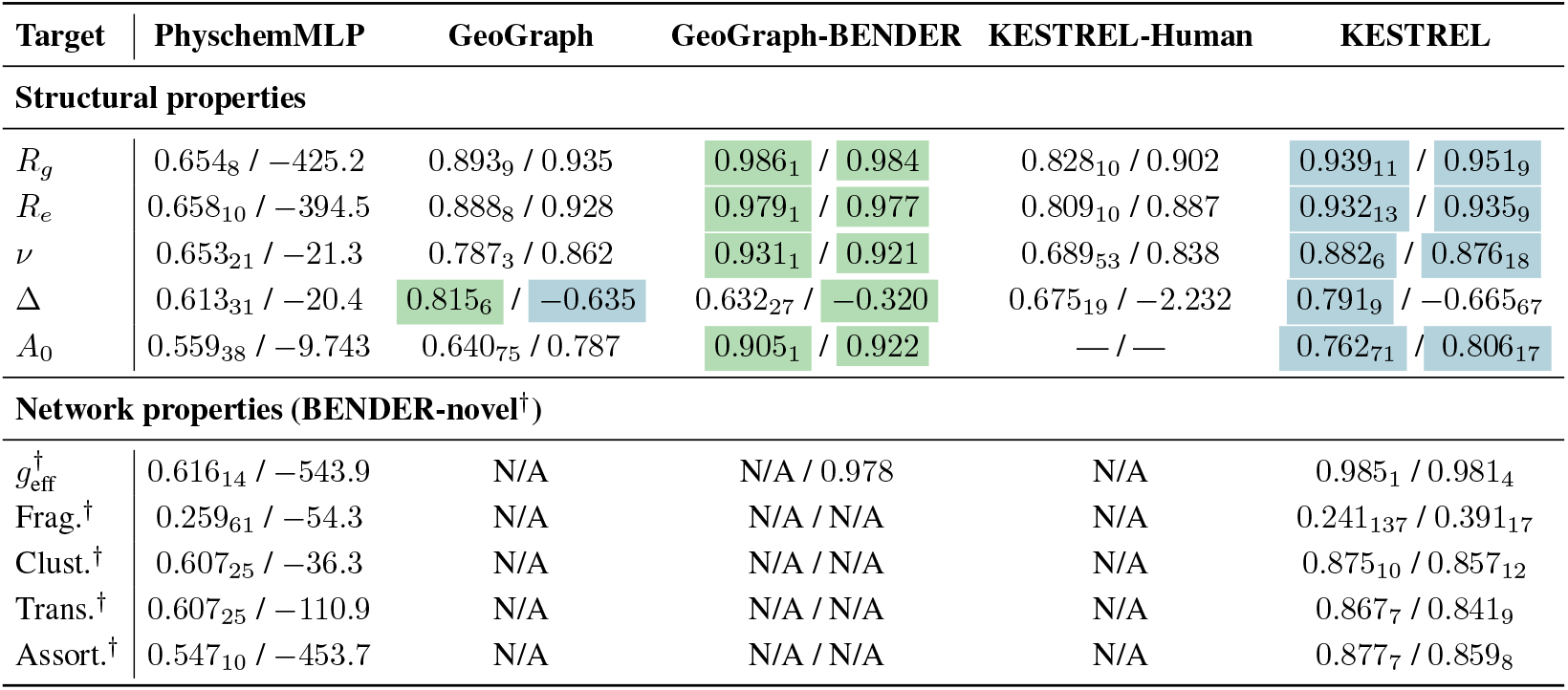
Benchmark *R*^2^ (test / OOD Virus) across all models and targets. N/A indicates the model does not predict that target. Subscripts denote standard deviations across 3 seeds, scaled by 10^−3^. Network property targets (†) are introduced by BENDER; no prior baselines exist. Best and second-best highlighted green and teal. KESTREL-Human is trained on Human-IDRome; its test column shows cross-distribution evaluation on the BENDER test split.

### 4.1 Cross-taxon training improves out-of-distribution generalisation

Table 2 decomposes OOD viral *ν* performance by architecture and training distribution, showing that including sequences from across the tree of life improves prediction in both architectures, with GeoGraph-BENDER achieving the highest *ν R*^2^ on this benchmark (0.921) despite training on approximately one-third the data.

**Table 2:** Architecture × data decomposition on OOD viral *ν*.

| $\nu$ OOD Virus | Human-only | Cross-taxon | gap |
| --- | --- | --- | --- |
| GeoGraph | 0.862 | **0.921** | +0.059 |
| KESTREL | 0.838 | 0.876 | +0.038 |

Table 3 isolates training distribution with everything else held constant, revealing that KESTREL-Human performs worse on all three geometric targets, with the *ν* gap (+0.038; seed SDs 0.030 and 0.017) comparable to the weaker arm’s seed spread.

**Table 3:**
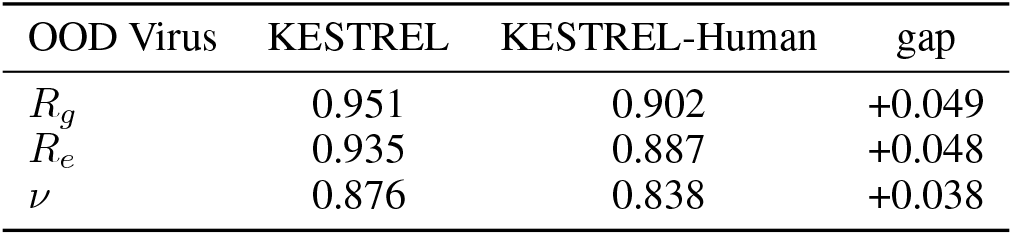
KESTREL vs KESTREL-Human on 1,025 OOD viral sequences (geometric targets).

| OOD Virus | KESTREL | KESTREL-Human | gap |
| --- | --- | --- | --- |
| $R_g$ | 0.951 | 0.902 | +0.049 |
| $R_e$ | 0.935 | 0.887 | +0.048 |
| $\nu$ | 0.876 | 0.838 | +0.038 |

On Human-IDRome, PhyschemMLP outperforms the transformer on *ν* (0.781 vs 0.694). On BENDER the ordering reverses (0.653 vs 0.882), and on OOD Viruses the compositional baseline collapses entirely (−21.3 vs 0.876).

### 4.2 Contact-network topology is conserved and sequence-predictable

Contact-network global efficiency is predicted with *R*^2^ = 0.981 on OOD Viruses (Figure 3), with clustering, transitivity, and assortativity showing similarly strong generalisation (*R*^2^ *>* 0.78). Per-taxon breakdown (Supplementary Table S1) shows *g*_eff_ ranging from 0.967 to 0.985 across all eleven evaluable taxa, including those with as few as 4 and 9 evaluation sequences.

A length-only baseline achieves *R*^2^ = 0.537 for *g*_eff_, while KESTREL reaches 0.963 on the length-residualised target, confirming that sequence context beyond chain length accounts for the majority of predictive accuracy. The no-embedding variant KESTREL-NoEmb achieves 0.977 versus 0.981 for the full model, indicating that learned residue representations contribute only marginally. Positive degree assortativity is observed across all 13 taxonomic groups, concentrated in [0.2, 0.4] with no taxon showing a negative median.

## 5 Discussion

The 2×2 decomposition in Table 2 provides a clean test of the cross-taxon diversity effect. Including sequences from across the tree of life improves out-of-distribution performance in both architectures tested, with the larger gain in GeoGraph (+0.059 versus +0.038 for KESTREL). This improvement cannot be explained by dataset size, as GeoGraph-BENDER trains on one-third the sequences of its Human-IDRome counterpart, nor is it architecture-specific, as the PhyschemMLP result makes plain. On human-only data, a 20k-parameter compositional model outperforms the 2.2M-parameter transformer, confirming that human IDP geometry is substantially determined by amino acid composition alone. On cross-taxon data the ordering reverses, and on held-out viral sequences it inverts entirely, with the compositional baseline collapsing to *R*^2^=−21.3 while KESTREL achieves 0.876. Architecture contributes only once the training data contains positional structure for it to exploit, and the diversity effect is therefore fundamentally attributable to the data.

The KESTREL versus KESTREL-Human *ν* gap (+0.038) is comparable in magnitude to the weaker arm’s seed spread, and we report it as directional rather than definitively established. The pattern is nonetheless consistent across all three geometric targets (*R_g_* +0.049, *R_e_* +0.048, *ν* +0.038), and the GeoGraph arm provides independent confirmation at a larger effect size using a fully distinct architecture and training regime.

Positive degree assortativity is consistent across all 13 taxonomic groups, and global efficiency is predicted with high accuracy on both held-out taxa. While this pattern is suggestive of a shared physical constraint, we note that *g*_eff_ has high mutual information with sequence length, which is itself similarly distributed across taxa, so the cross-taxon consistency may partly reflect shared length distributions rather than an independent topological law. That the majority of this predictability arises from sequence context beyond chain length indicates that the sequence-ensemble mapping governing contact-network architecture is as genuinely sequence-encoded as the geometric properties that accompany it. Whether this conservation reflects shared physicochemical constraints on disorder or convergent evolutionary pressures is a question BENDER opens rather than resolves.

The cross-taxon scope of BENDER’s condensate-relevant features makes it the first resource for studying how phase-separation propensity varies with evolutionary context at dataset scale, a question the human proteome alone cannot answer [21, 22, 15]. Saturation concentration and partition coefficient are fundamentally multi-chain quantities outside the scope of single-chain simulation, and extending BENDER to include slab-geometry simulations is the natural next step.

Although cross-force-field validation reveals no systematic *ν* bias across taxa, CALVADOS-2 is parameterised on eukaryotic experimental data and approximately 80% of per-sequence disagreement between the two force fields reflects genuine physical differences rather than pipeline noise, a limitation we state rather than explain away.

## 6 Conclusion

BENDER provides the first systematic test of whether coarse-grained IDP ensembles and the sequence–ensemble relationships they support generalise beyond the human proteome. CALVADOS-2 ensembles on taxonomically distant sequences prove no less reliable than on the eukaryotic data underlying force-field development, cross-taxon training consistently improves out-of-distribution prediction across two architecturally independent models with the benefit traceable to the data rather than model design, and contact-network topology is conserved and accurately sequence-predictable across all 13 taxonomic groups. Together, these results establish that the cross-taxon gap in IDP simulation was not merely a gap in coverage but a gap in understanding, and that the sequence–ensemble laws governing disordered chain behaviour are substantially conserved across deep evolutionary time in ways that become accessible only once training data spans it.

## Acknowledgment

All simulations were performed on the USF GAIVI cluster. We would like to thank Dr. Vladimir Uversky for his expert guidance and guidance on coarse-grained simulation protocol.

### A Species Breakdown

**Figure S1:**
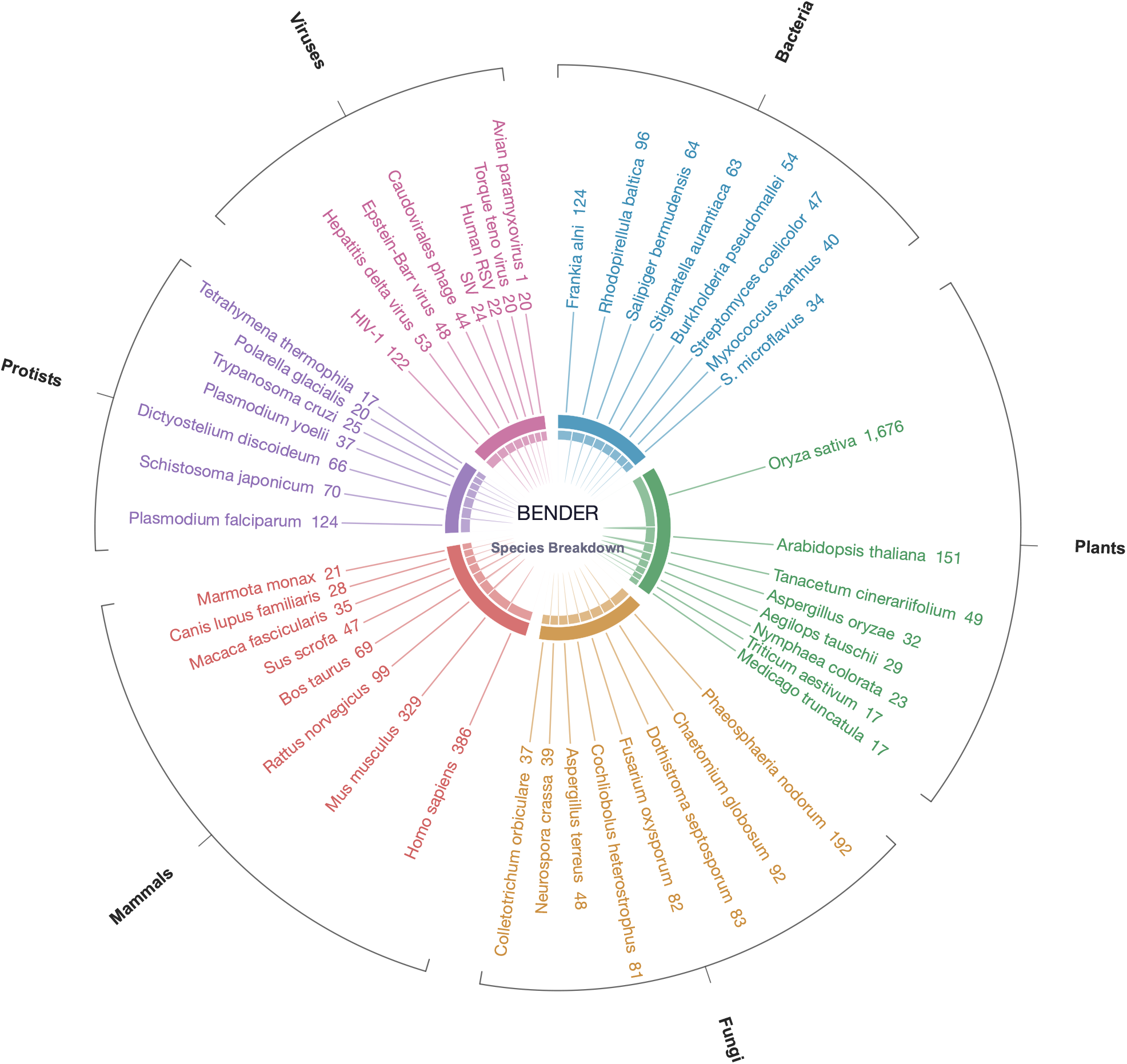
Species-level composition of BENDER, showing the top contributing organisms within each taxonomic group. Numbers indicate sequence counts per species.

### B KESTREL Architecture and Training

#### B.1 Architecture

KESTREL is a 2.2M-parameter transformer trained from scratch on one-hot encoded amino acid sequences. The backbone is a 4-layer Pre-Layer Normalization transformer encoder with hidden dimension 256, 8 attention heads, and feed-forward dimension 512, followed by mean pooling over the sequence length. A 16-dimensional taxon embedding is concatenated to the resulting sequence-level representation and passed to two shallow MLP prediction heads:

- **Geometric head**: predicts *R_g_*, *R_e_*, *ν*, Δ, and *A*_0_ (5 targets).
- **Network head**: predicts *g*_eff_, fragmentation index, average clustering coefficient, transitivity, and degree assortativity (5 targets).

Each head consists of two linear layers with GELU activation and dropout. The geometric and network heads are trained jointly with equal loss weighting.

#### B.2 Training Protocol

Training uses AdamW [31] with an initial learning rate of 1 × 10^−3^, cosine-annealed to 1 × 10^−5^ over the training run. Batch size is 64. Smoothed early stopping monitors validation loss with a patience of 20 epochs and a smoothing window of 5 epochs. Data is split into training (80%), validation (10%), and test (10%) using a cluster-aware partition based on CD-HIT clusters at 40% identity, ensuring that no cluster is split across partitions. Archaea and Viruses are held out entirely from training and used exclusively for OOD evaluation.

All results are reported as mean ± SD across 3 random seeds (42, 67, 93). The subscripts in Table 1 denote seed SDs scaled by 10^−3^.

#### B.3 Taxon Embedding

A 16-dimensional learned taxon embedding is concatenated to the sequence representation before the prediction heads. For OOD taxa (Viruses and Archaea), a zero vector is used at inference. To verify this does not inflate OOD performance, we train KESTREL-NoEmb, a variant with the taxon embedding removed entirely. On OOD Viral *g*_eff_, KESTREL-NoEmb achieves *R*^2^ = 0.977 versus 0.981 for the full model, confirming the zero-vector strategy does not artificially boost performance.

#### B.4 KESTREL-Human

KESTREL-Human is an architecture-matched variant without the taxon embedding, trained on Human-IDRome [9] using the same similarity-based split as GeoGraph [8]. The training protocol (optimiser, learning rate schedule, early stopping) is identical to KESTREL. This ensures that differences in OOD performance between KESTREL and KESTREL-Human are attributable solely to the training distribution, not to architectural or optimisation differences.

### C PhyschemMLP

PhyschemMLP is a three-layer MLP with approximately 20k parameters mapping 13 scalar physicochemical descriptors directly to all 10 prediction targets, with no positional sequence information. The input features are:

1. Fraction of charged residues (FCR)
2. Net charge per residue (NCPR)
3. Fraction of positive residues (*f*_+_)
4. Fraction of negative residues (*f_−_*)
5. Fraction of aromatic residues (*f*_aro_)
6. Fraction of hydrophobic residues (*f*_hydro_)
7. Fraction of polar residues (*f*_polar_)
8. Fraction of disorder-promoting residues (*f*_dis_)
9. Fraction of order-promoting residues (*f*_ord_)
10. Fraction of proline (*f_P_*)
11. Fraction of glycine (*f_G_*)
12. Sequence charge decoration (SCD)
13. Sequence length (log-transformed)

The MLP uses hidden dimensions [128, 64, 32] with GELU activations and dropout of 0.1. It is trained on both BENDER and Human-IDRome under the same protocol as KESTREL (AdamW, cosine annealing, smoothed early stopping). Where PhyschemMLP matches or exceeds KESTREL, the target is composition-determined; where KESTREL substantially outperforms it, positional sequence context is essential.

### D GeoGraph Retraining

GeoGraph [8] is a graph neural network that combines ESM-2 embeddings with a residue-level contact-network architecture, trained on 28,058 Human-IDRome sequences with *g*_eff_ as an auxiliary target. It represents the current state of the art on geometric IDP ensemble targets.

To produce GeoGraph-BENDER, we retrained GeoGraph end-to-end on the BENDER training split using its published protocol and hyperparameters without modification. The same cluster-aware 80/10/10 split and OOD holdout design used for KESTREL were applied. This ensures that the comparison between GeoGraph (IDRome-trained) and GeoGraph-BENDER isolates the effect of training data within a single architecture.

We note that GeoGraph’s hyperparameters were originally tuned on Human-IDRome and were not retuned for BENDER. Despite this potential disadvantage, GeoGraph-BENDER achieves *ν R*^2^ = 0.921 on OOD Viruses, surpassing both its IDRome-trained version (0.862) and KESTREL (0.876).

### E Sequence Leakage Analysis

To verify that OOD performance is not inflated by sequence similarity to training data, we performed a BLAST search of all OOD sequences against the training set. Only 1 of 1,025 OOD Virus sequences has training homologs above 50% identity at ≥80% coverage. All three are short conserved peptides under 65 residues. Removing these sequences does not change any reported *R*^2^ value by more than 0.001.

The cluster-aware partition ensures that sequences within the same CD-HIT 40% identity cluster are never split across training and evaluation sets, providing a stricter guarantee against leakage than random splitting.

### F Convergence Quality

Convergence of the CALVADOS-2 simulations was assessed using the *R*^2^ of the power-law fit to internal scaling distances (nu_fit_r2). Across all 11,533 sequences, the median fit *R*^2^ = 0.998, with 95.2% of sequences exceeding 0.95 and 87.4% exceeding 0.99. The median per-sequence *ν* uncertainty from the fit covariance is 0.0025, corresponding to 3.9% of the population standard deviation. The per-taxon distribution is shown in Figure 4 of the main text.

### G Feature Extraction Details

#### G.1 Geometric Properties

**Radius of gyration (***R_g_*). Computed as the mass-weighted root-mean-square distance of all beads from their centre of mass, averaged over production frames.

**End-to-end distance (***R_e_*). Euclidean distance between the first and last *C_α_* beads, averaged over production frames.

**Flory scaling exponent (***ν***) and prefactor (***A*_0_). Fitted from a power law 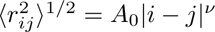 between inter-residue distances and sequence separation, excluding pairs separated by fewer than 5 residues. This convention matches Human-IDRome [9] and GeoGraph [8].

**Asphericity (**Δ). Computed from the eigenvalues *λ*_1_ ≤ *λ*_2_ ≤ *λ*_3_ of the gyration tensor:

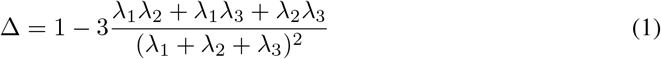

Values range from 0 (sphere) to 1 (rod), averaged over production frames.

#### G.2 Contact-Network Properties

The *C_α_*–*C_α_* contact graph is constructed for each frame using a 0.8 nm cutoff. Five graph topology metrics are computed from the ensemble-averaged contact frequency matrix.

**Global efficiency (***g***_eff_).** Average inverse shortest path length across all node pairs, quantifying how efficiently conformational signals propagate across the chain [28].

**Fragmentation index.** Fraction of residues in the largest connected component, quantifying chain connectivity and multivalent interaction capacity.

**Average clustering coefficient.** Mean local clustering across all residues, capturing locally dense contact neighbourhoods associated with transient secondary structure.

**Transitivity.** Global prevalence of closed contact triplets, complementing clustering as a measure of network cohesion.

**Degree assortativity.** Pearson correlation of node degrees across edges, measuring whether highly connected residues preferentially contact other highly connected residues [29, 30].

#### G.3 Phase-Separation-Relevant Features

Per-sequence pi-pi and cation-pi contact frequencies are computed from the simulation trajectories by counting pairwise contacts between aromatic residues (Phe, Tyr, Trp) and between cationic residues (Arg, Lys) and aromatic residues, respectively, normalised by sequence length and number of production frames.

### H Per-Taxon Benchmark Results

**Table S1:**
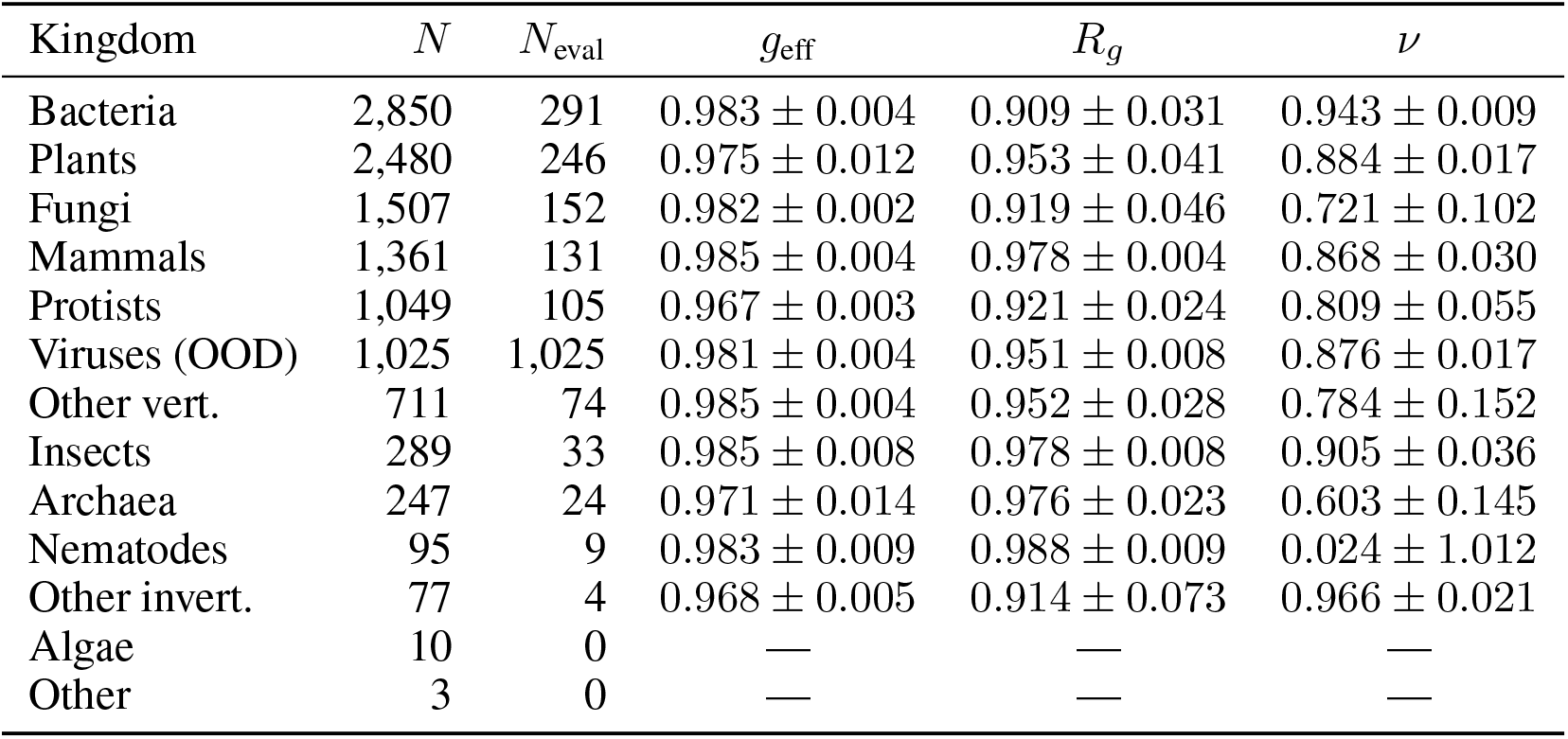
Per-taxon *R*^2^ (mean ± SD, 3 seeds) on the expanded dataset (*n* = 11,704), Virus-held-out run.

### I Per-Taxon Target Distributions

**Figure S1:**
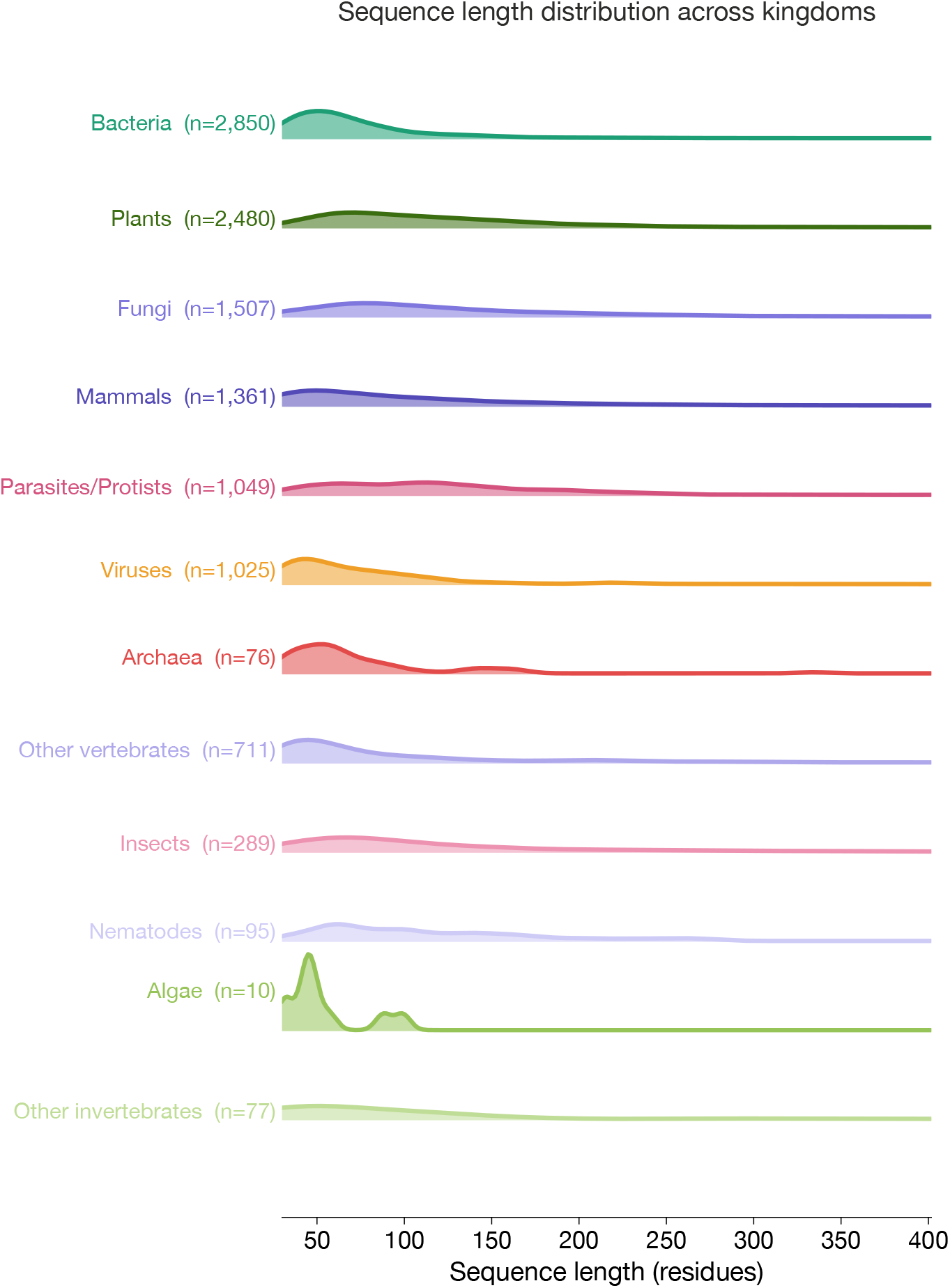
Sequence length distribution across taxa. Ridge density plots of IDP sequence lengths (residues) across 12 taxonomic taxa. Bacteria show a sharply defined peak relative to Plants or Mammals. Viruses show a distribution similar to Bacteria, consistent with compact viral proteomes. Protists display a longer-tailed, bimodal distribution.

**Figure S2:**
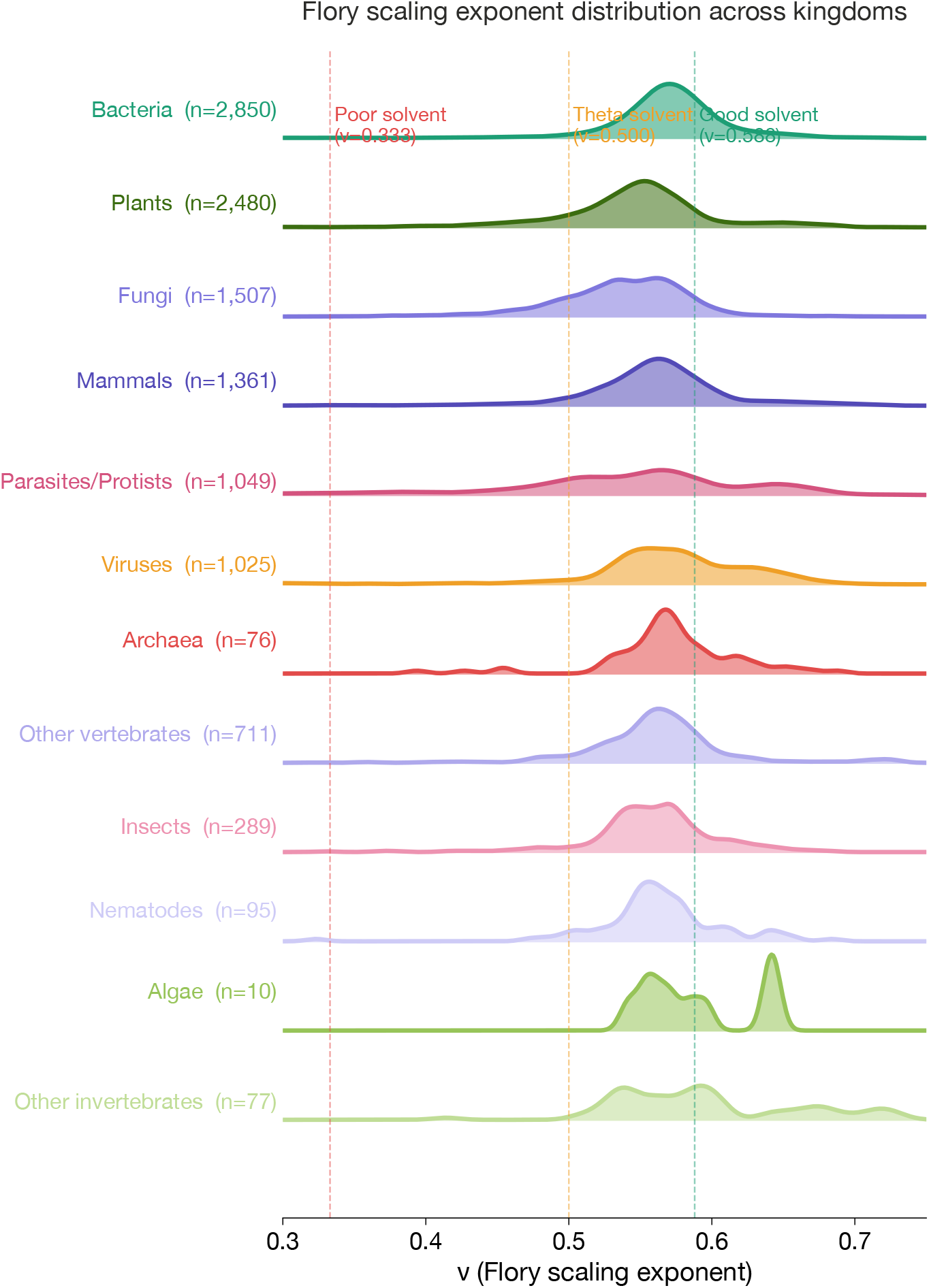
Flory scaling exponent distribution across taxa. Dashed vertical lines indicate canonical polymer physics regimes: good solvent (*ν* = 0.588), theta solvent (*ν* = 0.500), and poor solvent (*ν* = 0.333). Bacteria show the broadest distribution. Mammals show a tight distribution near *ν* ≈ 0.57, consistent with the human IDRome [9]. Protists and Viruses show wider distributions with substantial density above the good solvent limit. BENDER collectively spans the full physically accessible range of *ν*, substantially broader than human-centric datasets.

**Figure S3:**
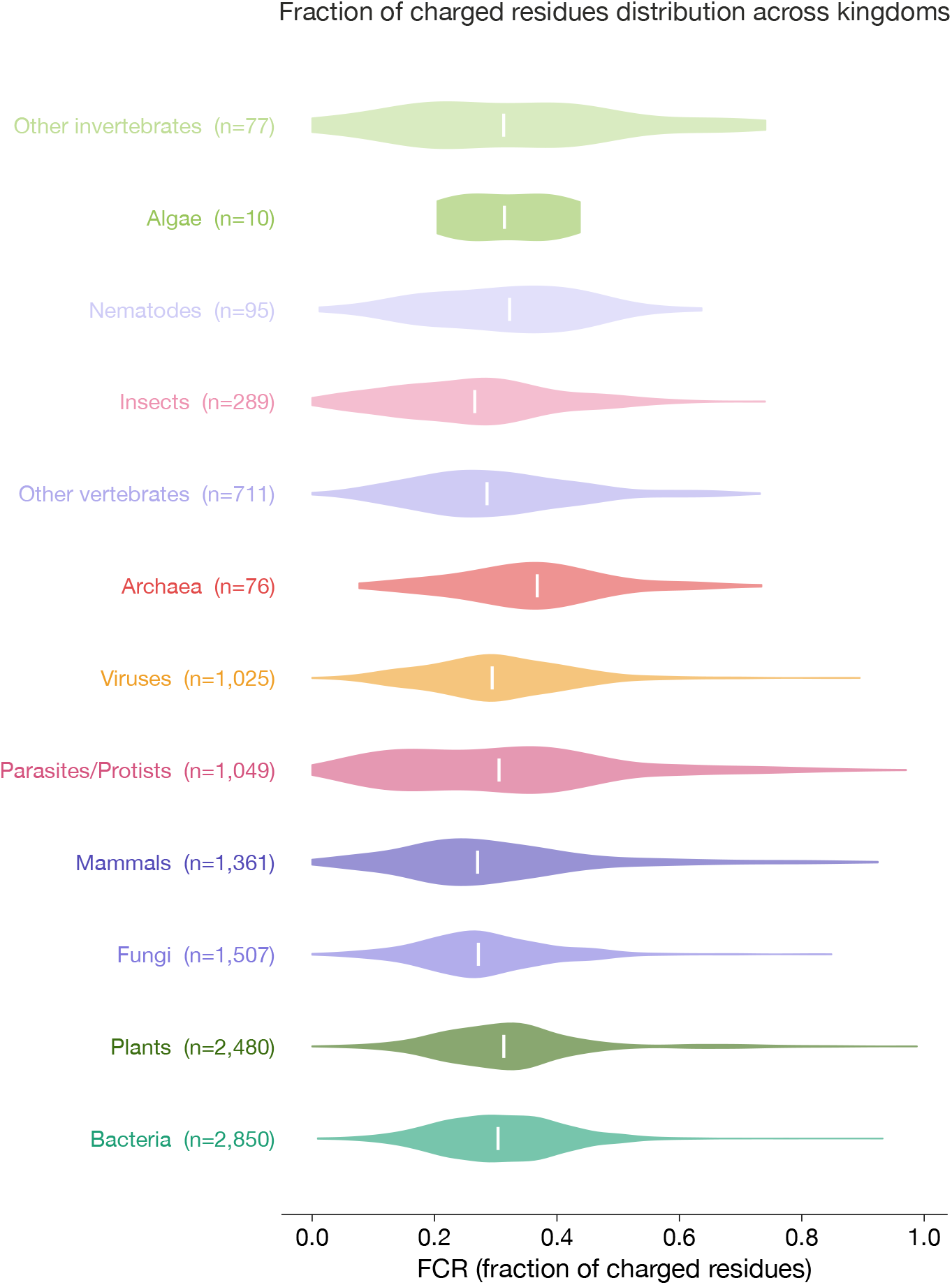
Fraction of charged residues (FCR) across taxa. All taxa show similar median FCR near 0.25–0.30 but differ in distributional shape. Cross-taxa conservation near 0.25–0.30 suggests a shared functional constraint on charged residue content in disordered proteins.

**Figure S4:**
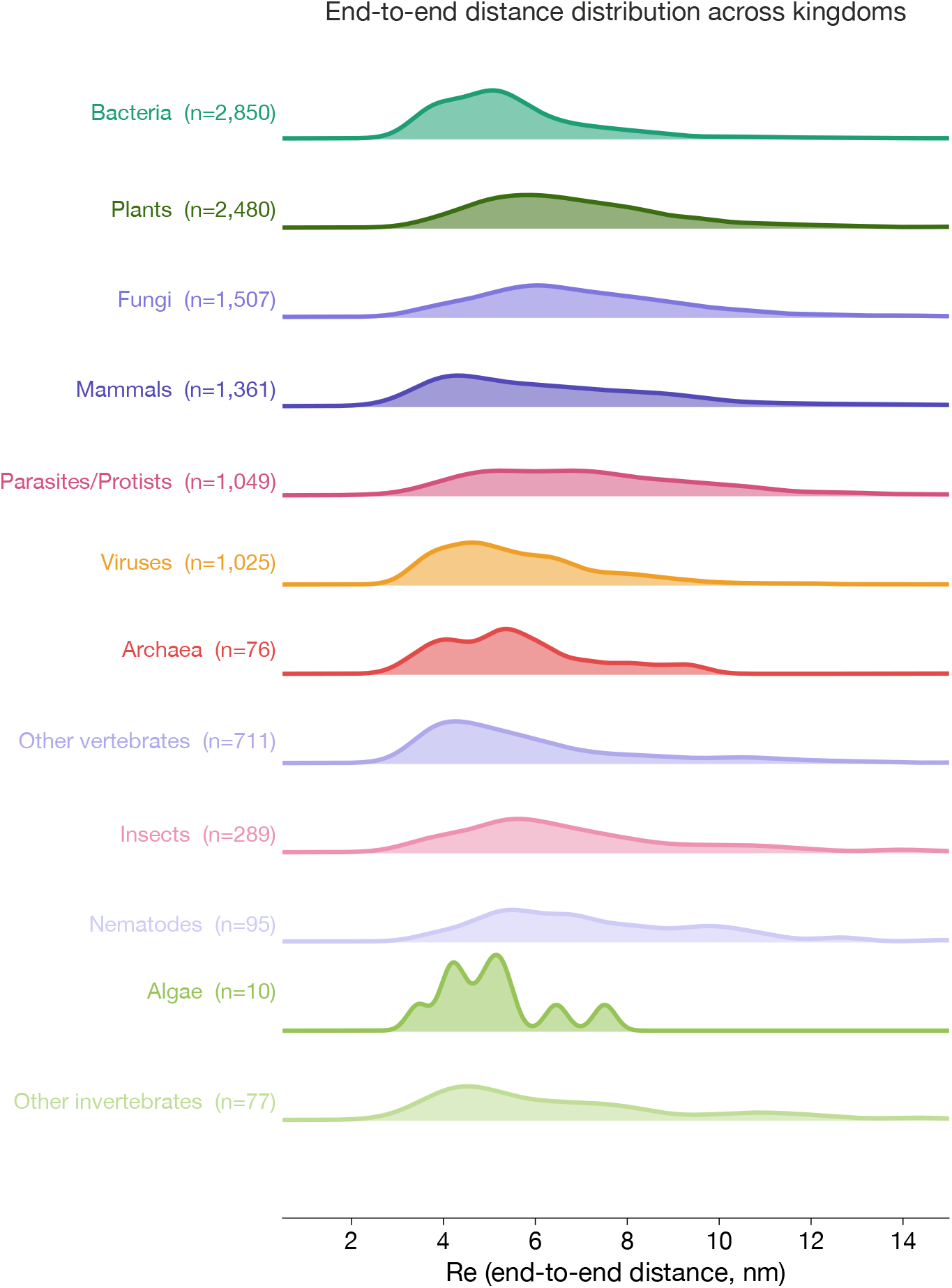
End-to-end distance (*R_e_*) distribution across taxa. Bacteria show a sharp peak near 3–4 nm. Plants show a broader distribution extending to larger values. Archaea display a bimodal shape reflecting two distinct length populations.

**Figure S5:**
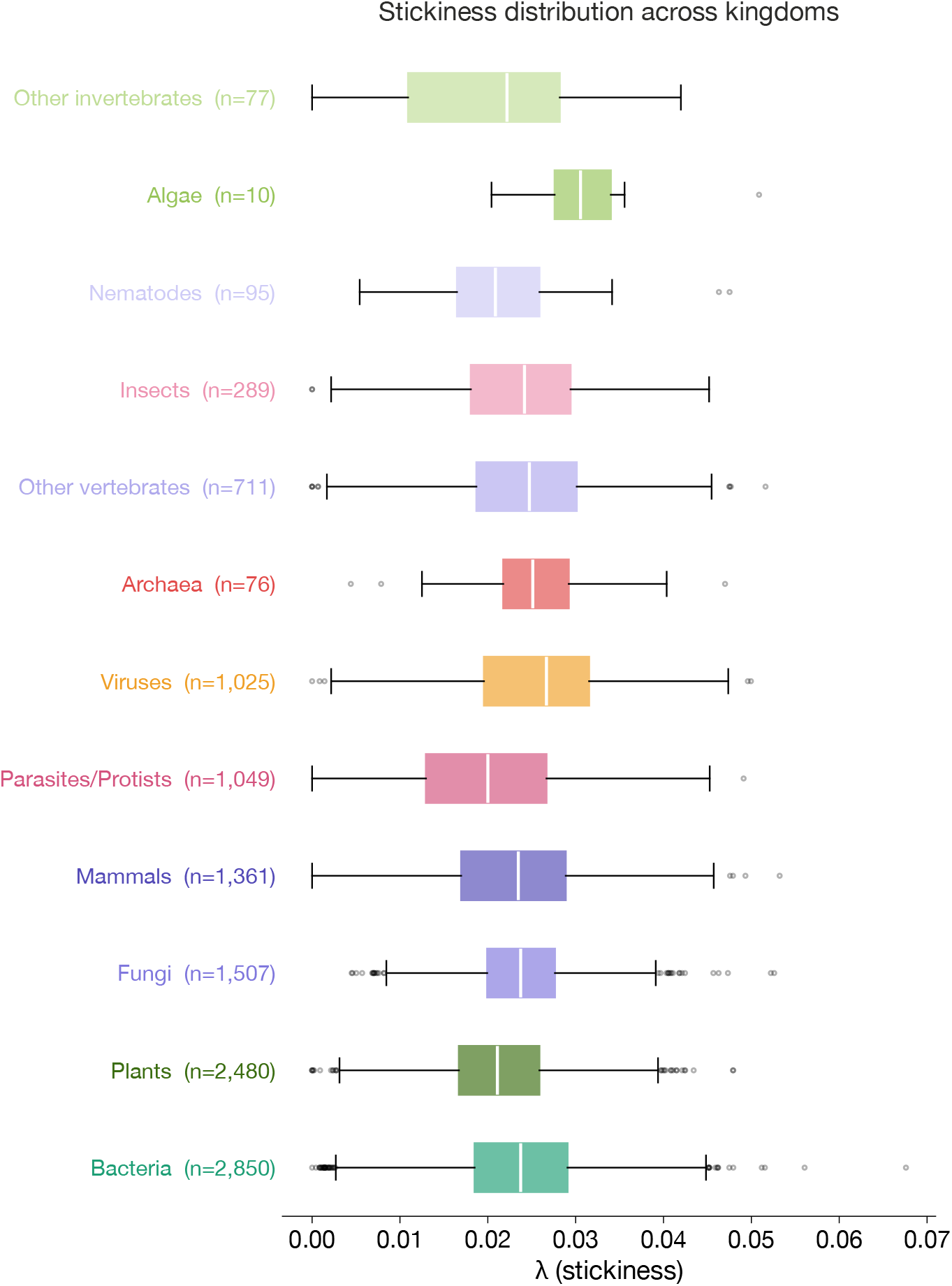
Stickiness (*λ*) distribution across taxa. Values are consistently low across all taxa (median ≈ 0.02–0.03), reflecting the known depletion of hydrophobic residues in fully disordered sequences. Viruses show slightly elevated stickiness, consistent with aromatic enrichment in viral disordered regions.

**Figure S6:**
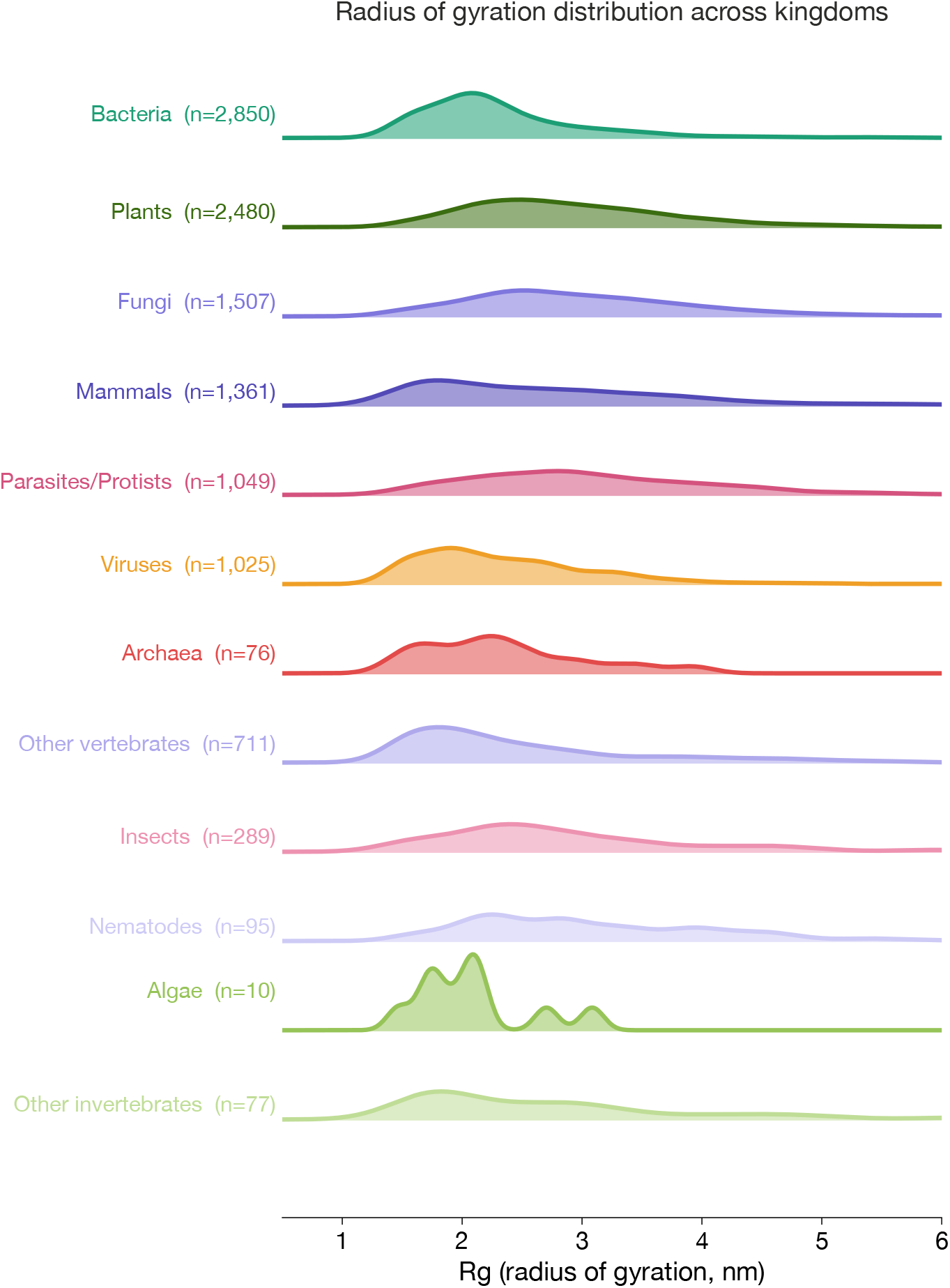
Radius of gyration (*R_g_*) distribution across taxa. Bacteria and Plants show the broadest distributions with extended right tails. Mammals show a tight peak near 1.5 nm. Archaea show a bimodal distribution consistent with the two length populations observed in Figure S1.

**Figure S7:**
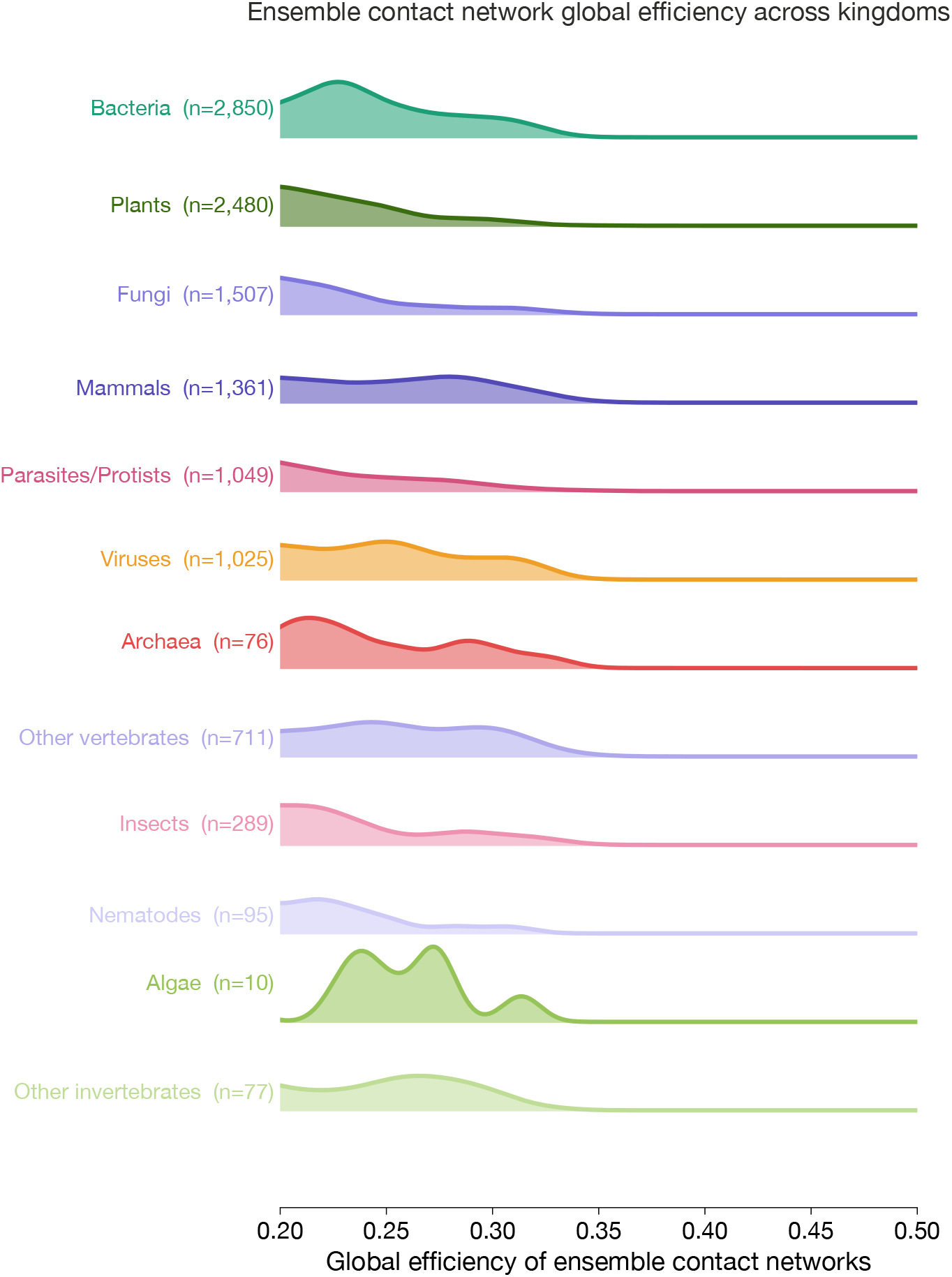
Global efficiency (*g*_eff_) distribution across taxa. All taxa show distributions concentrated in 0.2–0.4 with peaks near 0.25–0.30. Viruses and Archaea show slightly broader distributions, supporting their use as OOD evaluation taxa.

**Figure S8:**
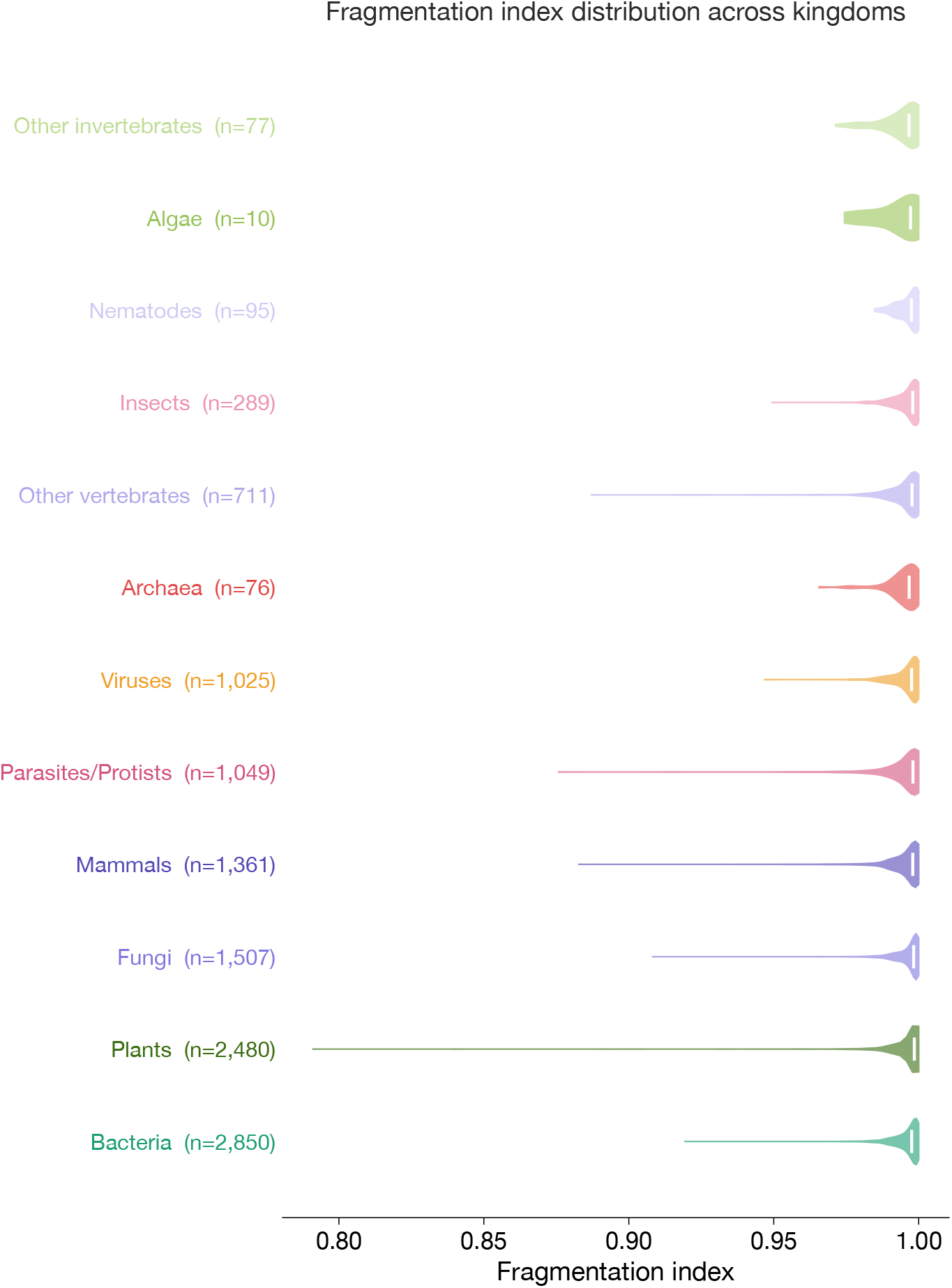
Fragmentation index distribution across taxa. All taxa show fragmentation indices concentrated near 1.0, indicating largely cohesive contact networks. Plants show the broadest distribution with the longest left tail.

**Figure S9:**
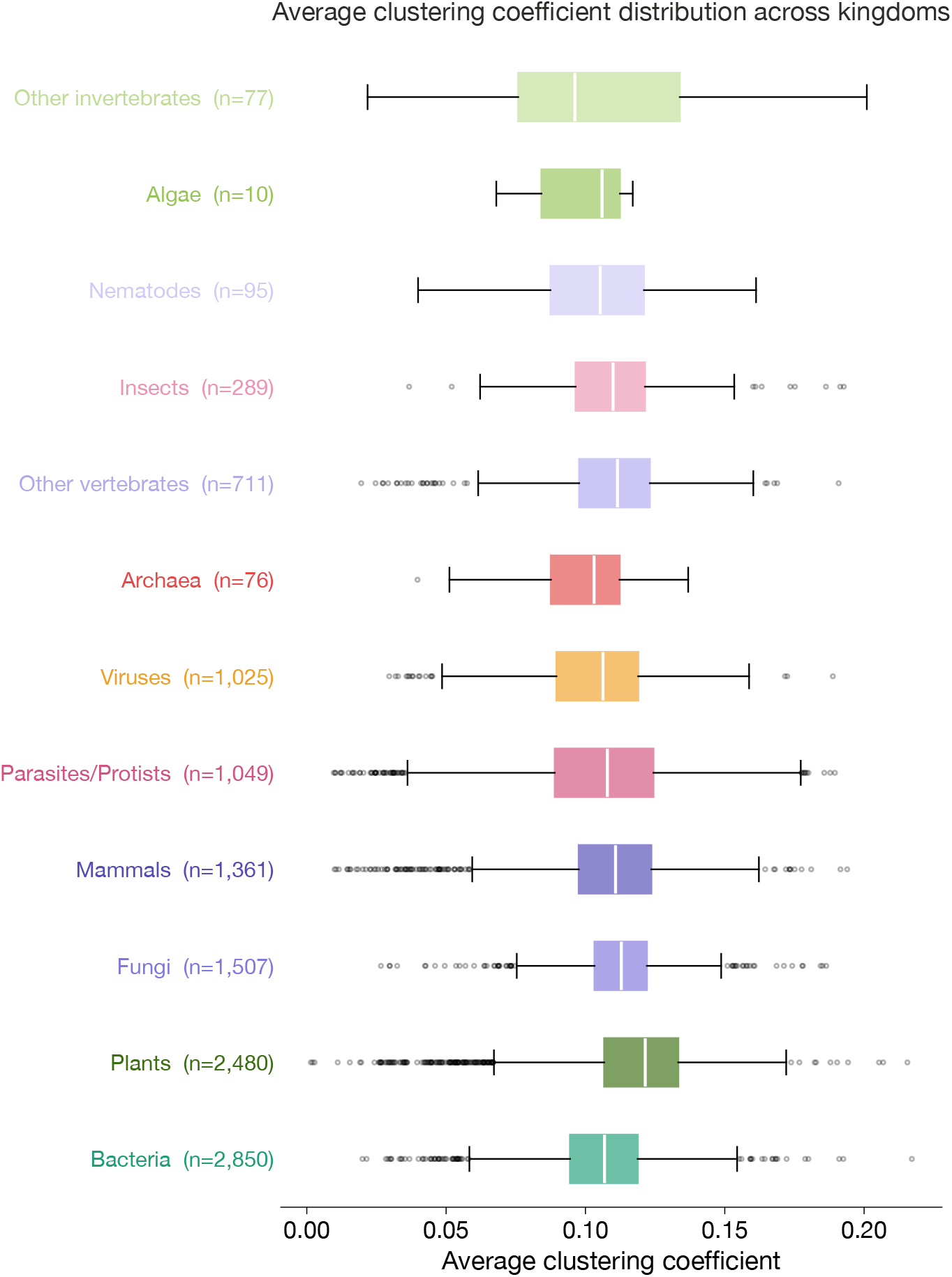
Average clustering coefficient distribution across taxa. Values are consistently low (0.05–0.15) across all taxa, reflecting the sparse, chain-like topology of IDP contact networks. All taxa share similar median clustering near 0.10.

**Figure S10:**
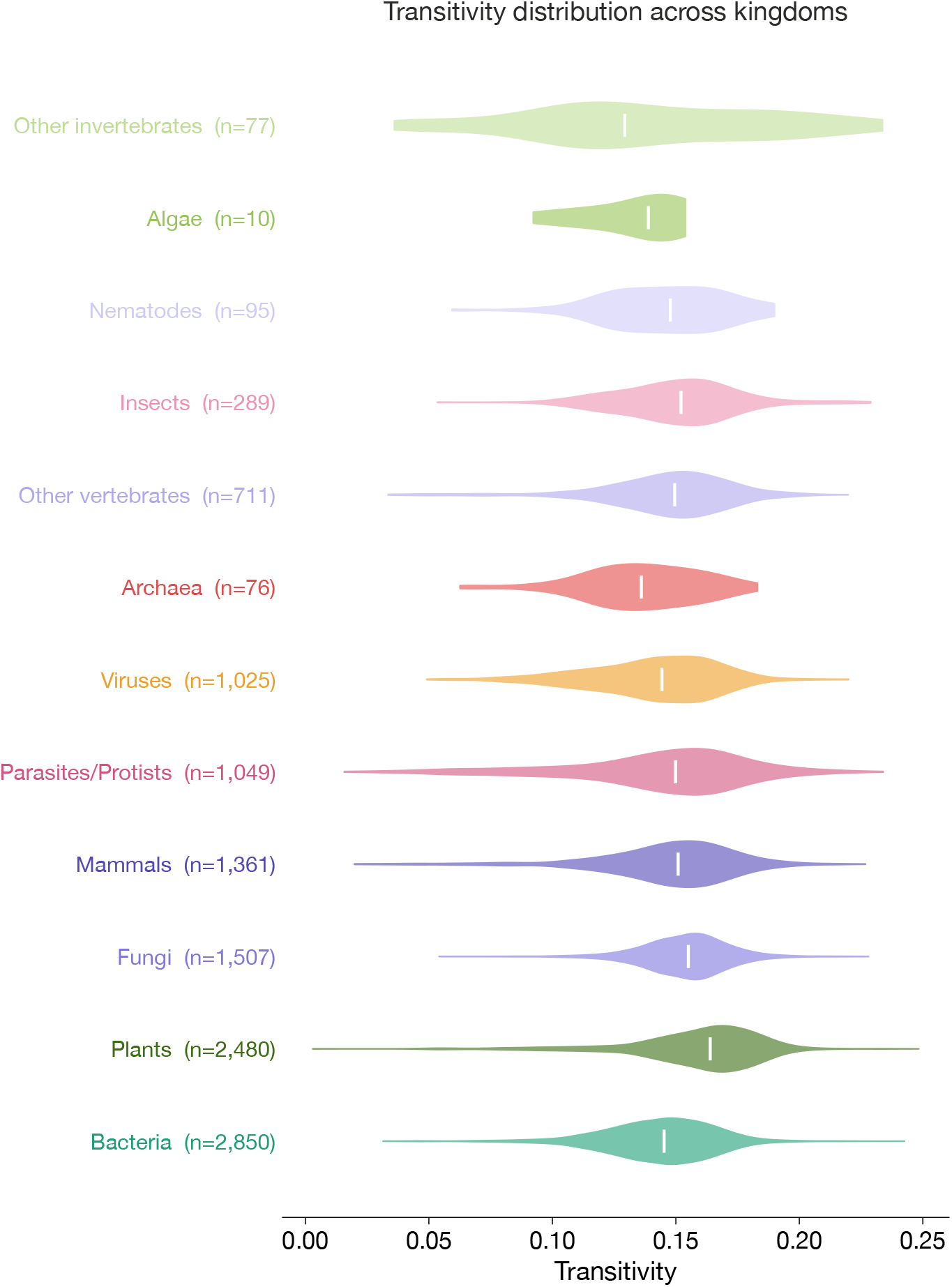
Transitivity distribution across taxa. Values concentrated in 0.10–0.20, closely mirroring the clustering coefficient distributions. Viruses and Archaea show slightly lower median transitivity relative to eukaryotic taxa.

**Figure S11:**
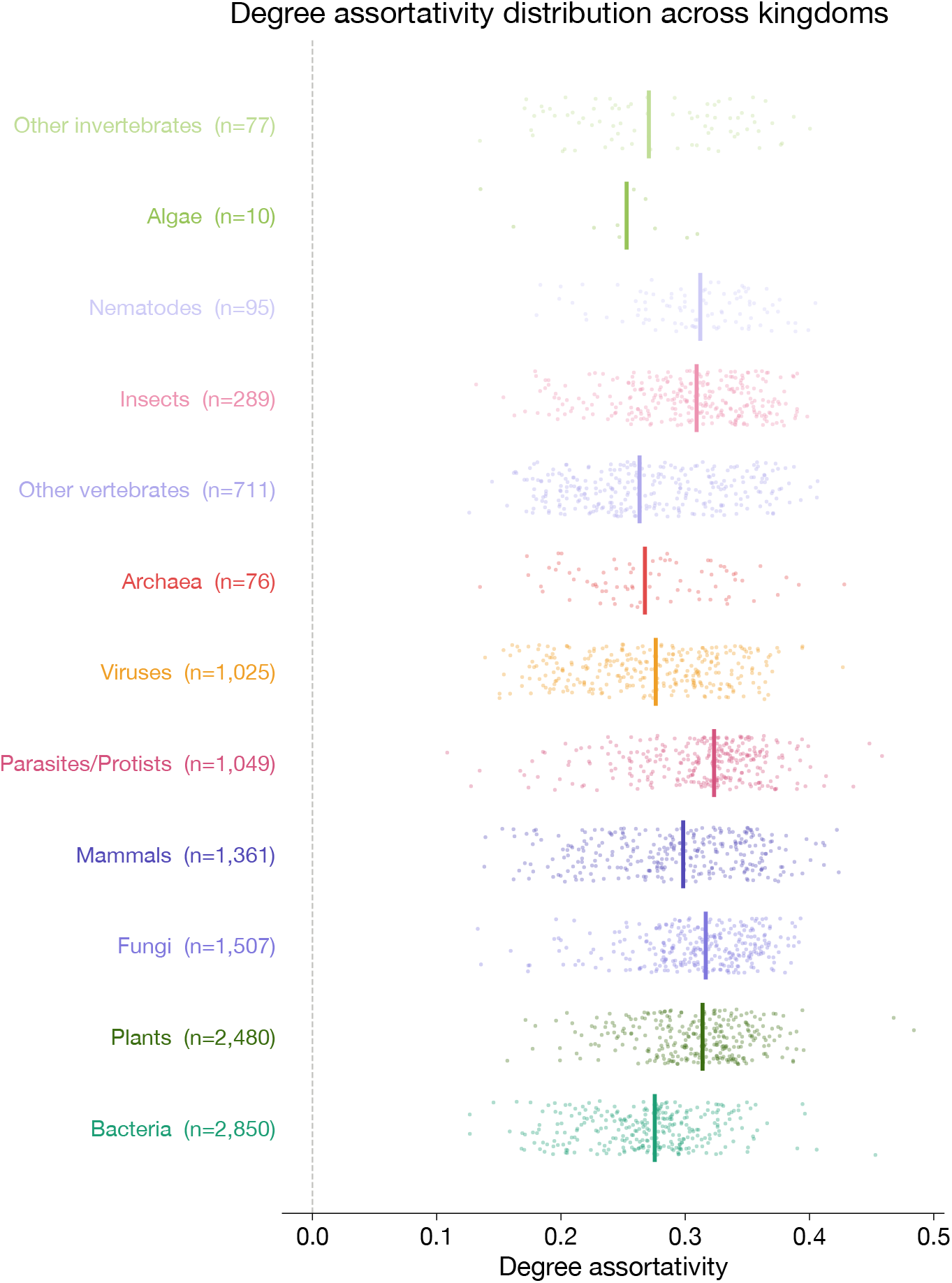
Degree assortativity distribution across taxa. All taxa show positive degree assortativity concentrated in 0.2–0.4. No taxon shows a negative median. The tight cross-taxa conservation, despite billions of years of independent evolution, suggests that distributed hub topology is a conserved physical feature of disordered protein contact networks.

## Notes

### Competing Interest Statement

The authors have declared no competing interest.

### Summary of Updates

The tile was updated along with the focus of the paper, clarifying the strength and application areas of the BENDER dataset discussed.

